# 3D-genome impacts of TAD-boundary-disrupting ultra-rare structural variants identified in schizophrenia

**DOI:** 10.64898/2026.10.26.761993

**Authors:** Luz M. Porras, Martilias Farrell, Isabelle Rodriguez-Lausell, Gabriel A. Iglesias-Maldonado, Emily Val F. Tuliao, Elizabeth Wright, Amphitreatte Kreiman, NaEshia Ancalade, Christopher Stoddard, Jin P. Szatkiewicz, Paola Giusti-Rodríguez

**Author notes:** Co-senior/co-corresponding authors: Paola Giusti-Rodríguez, PhD. University of Florida College of Medicine, Department of Psychiatry, 1149 Newell Dr, L4-100D, Gainesville, FL 32610., Jin Szatkiewicz, PhD. University of North Carolina at Chapel Hill, Department of Genetics, 5097 Genetic Medicine Building, CB# 7264, Chapel Hill, NC 27599.

## Abstract

Schizophrenia is a complex and multifactorial brain disorder that impacts ∼1% of the U.S. population. The genetic architecture of schizophrenia includes 10 large rare structural variants (SVs) ∼10 exonic variants, and over 250 common variants. Previously we combined whole genome sequencing (WGS) with brain-specific epigenomic data and found an excess of ultrarare SVs in schizophrenia localizing to the boundaries of topologically associated domains (TADs). Alterations in TAD boundaries (TAD-b) may lead to disarrangement of enhancer-promoter contacts and dysregulation of gene expression and have been associated with several human diseases. Our study sought to functionally characterize two of these TAD-b SVs using chromatin conformation capture (e.g., Hi-C) and RNA sequencing, to unveil mechanistic insight into the effects of these variants on 3D-genome organization and gene expression. To achieve this, we genetically engineered these two TAD-b SVs in isogenic induced pluripotent stem cells (iPSCs) using CRISPR/Cas9. iPSCs were differentiated into neuronal precursor cells, excitatory glutamatergic neurons, and medium spiny neurons. We found that these TAD-boundary-disrupting SVs could reshape the 3D-genome and transcriptional networks in a cell-type-specific manner.

## Introduction

Schizophrenia is a highly heritable psychiatric disorder whose genetic architecture includes both common and rare genetic variation (Owen et al. 2023). Despite substantial progress in gene discovery, the mechanisms by which many risk variants contribute to disease remain poorly understood, particularly in the case of structural variants (SVs). SVs can contribute to disease risk through both coding and regulatory mechanisms (Marshall et al. 2017) and can exert direct pathogenic effects, as exemplified by intragenic deletions in *ABCB11,* a bile acid transporter implicated in antipsychotic response, found in treatment-resistant schizophrenia and enriched in neurons of the mesocortical and mesolimbic dopamine pathways (Maury et al. 2023). They can also act indirectly by reshaping gene regulation without altering coding sequences. For instance, 5′-UTR deletions in *NRXN1* generate ectopic, allele-specific chromatin loops that connect cryptic promoters to distal enhancers, dysregulating transcription through altered 3D genome architecture (Maury et al. 2023). These findings highlight a broader principle: SVs often produce pathogenic effects via 3D genome dysregulation, frequently manifesting as classic position effects (Sreenivasan et al. 2025). Consistent with this model, we previously identified ultra-rare SVs in individuals with schizophrenia that overlapped topologically associated domain boundaries (TAD-b), which may lead to disrupted chromatin organization, and contribute to disease risk (Halvorsen et al. 2020).

Evidence from other biological contexts further strengthens the link between SVs and 3D genome disruption. In cancer, Li et al. identified 185 SVs that disrupt TAD-bs (TAD-SVs) in lymphoblastoid cell lines using an integrative Hi-C pipeline, many of which overlap candidate cis-regulatory elements (cCREs) (Li et al. 2024). Their analysis showed that SVs were not isolated abnormalities, but recurrent events that reorganized 3D-genome organization, rewired enhancer–promoter contacts, and misregulated gene expression (Li et al. 2024). Similar mechanisms have been established in developmental disorders, where high-resolution 3D genome mapping (Hi-C, promoter-capture Hi-C, Micro-C, ChIA-PET) has shown that that SVs can reposition enhancers, create ectopic loops, or remove boundary insulation, leading to pathogenic transcriptional programs(Lupiáñez et al. 2015; Halvorsen et al. 2020). Analytical tools such as nMAGMA extend these insights to complex traits by leveraging Hi-C contact maps to link noncoding variants to target genes via spatial proximity, revealing additional schizophrenia-relevant biological pathways (Yang et al. 2021).

The relevance of 3D-genome organization is particularly pronounced in the human brain. Chromatin architecture in the adult cortex spans from nucleosome-level interactions to megabase-scale domains (Hu et al. 2021; Lu et al. 2020; Giusti-Rodríguez et al. 2019). Large-scale epigenomic profiling of 739 prefrontal cortex samples identified thousands of cis-regulatory domains (CRDs) that align with Hi-C–defined TADs. Notably, clusters of hyper-acetylated CRDs were enriched for schizophrenia heritability and associated with fetal development and glutamatergic neuron signaling (Girdhar et al. 2022), underscoring the regulatory importance of chromatin topology in psychiatric disease.

Network-level studies of enhancer–promoter interactions (EPIs) reveal that regulatory architecture is structured and disease-relevant, with neuropsychiatric disorder–associated variants enriched in specific cell types and network modules (Koesterich et al. 2024). These networks are also temporally dynamic during neural differentiation, showing stage-specific variant enrichments (DeGroat et al. 2024), highlighting the need to resolve regulatory architecture by cell type and developmental stage.

Single-cell studies further demonstrate that genetic and epigenetic perturbations produce highly cell-type-specific effects. In the Setd1a haploinsufficiency model, transcriptional changes vary across neuronal populations, with Foxp2⁺ neurons being the most affected (Chen et al. 2022). Multi-omic analyses in human neurons and glia show similar specificity: schizophrenia risk maps to deeper-layer projection neurons, bipolar disorder to upper-layer neurons, and parvalbumin interneurons exhibit shared vulnerability across disorders (Hu et al. 2021). Together, these findings underscore that disease mechanisms cannot be captured by bulk tissue and require cell type specific resolution. Here, we aim to characterize the 3D-genome effects of ultra-rare SVs associated with schizophrenia, focusing on those that colocalize with TAD-b or nearby regions, and assessing their impact on local and genome-wide chromatin organization. We further investigate how these variants influence gene expression across the genome. Given the critical role of cell type-specific regulatory effects, we specifically examined developmentally relevant and schizophrenia-associated cell types, including excitatory glutamatergic neurons (EGN), medium spiny neurons (MSN), and neural precursor cells (NPC) (Bryois et al. 2020).

## Methods

### Rationale

Given our WGS findings, we hypothesized that TAD-b-disrupting SVs could be particularly relevant for the etiology of schizophrenia. To examine this, we used CRISPR-Cas9 to genetically engineer two TAD-boundary-disrupting deletions identified in our schizophrenia WGS cohort in isogenic human induced pluripotent stem cells (iPSCs), differentiated these into NPCs, EGNs, and MSNs, and characterized these using deep, genome-wide Hi-C and bulk RNA sequencing.

### Selection of TAD-boundary-disrupting SVs (TAD-b SVs) for modeling using CRISPR -Cas9

Since the TAD-b SVs were initially detected from WGS data, we first aimed to verify the presence of these variants using digital PCR. We attempted validation of 17 TAD-b deletions found in schizophrenia cases. For each deletion, we designed custom probes with three probes per variant placed near the start, end, and middle of the region of interest. A deletion was confirmed when its target probes had half the intensity of the control probes as measured by the Fam/Vic ratio. After confirming that all 17 TAD-b deletions, 2 loci were eventually selected for modeling with CRISPR/Cas9. Our prioritization strategies are briefly described here. First, we excluded deletions that directly interfered with a gene based on the hg38 genome build, due to the potential confounders that this could introduce to the experimental measurement of TAD-b disruption effects. Next, we annotated the remaining deletions using a variety of datasets and measures, including: the size of the deletion, whether the deletion affected brain specific insulators, the number of genes in the neighboring TADs but outside of the TAD-b, pLI score of these genes (i.e. intolerant to LoF variation) (Karczewski et al. 2020; Lek et al. 2016), whether these genes had an OMIM entry (Amberger et al. 2019), whether and when these genes were expressed in the brain, the number of unique eQTL-SNPs in any tissue (GTEx Consortium 2020), if this was a region previously associated with a psychiatric disorder or brain trait (Lu et al. 2020), conservation score based on data from the 200 Mammals Project (Sullivan et al. 2023), presence of known regulatory elements (based on brain open chromatin (Bryois et al. 2018), ChIP-seq (Lu et al. 2020), Hi-C data (Giusti-Rodríguez et al. 2019), TFBS (Castro-Mondragon et al. 2022), VISTA Enhancer Browser (Visel et al. 2007), Virtual 4C-seq (Wang et al. 2018); and location and orientation of CTCF motifs within the SV (Lu et al. 2020).

### Genetic engineering of SVs using CRISPR-Cas9

We used CRISPR/Cas9 to engineer hiPSC lines harboring the desired deletions. Two guide RNAs were chosen that flanked the region to be deleted using CRISPR RGEN tools (http://www.rgenome.net/). The guide RNAs were cloned into pSpCas9(BB)-2A-Puro (PX459) V2.0 (gift from Feng Zhang; RRID: Addgene_62988) after digestion with Bbs1 to allow for ligation of the guide RNA insert. Human iPSCs from Coriell Institute for Medical Research, specifically cell line GM25256 iPSCs from a healthy individual, were grown on mitotically inactivated MEFs. 24 hours prior to targeting, cells were treated with ROCKi (Y-27632). The following day, cells were dissociated with Accutase, pelleted, and resuspended according to the 4D nucleofection Primary P3 kit protocol (Lonza), before nucleofection using 2 µgs of each CRISPR plasmid and program CB-150. Cells were then plated on DR4 MEFs supplemented with ROCKi. The following day 0.5-1 ng/µl of puromycin and ROCKi was added to fresh media. This selection was continued for 48 hours total to select cells transiently expressing the vector containing the guide RNA and Cas9 protein; no integration of the CRISPR Cas9 was made into the genome. Cells were maintained in ROCKi-containing media until small colonies were observed. After this point, cell culture continued with daily fresh media changes. After a total of 15 days, colonies were isolated and plated into 24-well plates. Four days later, several colonies from each well were isolated, and DNA was extracted using the HOT SHOT method to prepare a template for PCR (Meeker et al. 2007).

### Confirmation of successful genetic engineering of selected deletions

PCR was performed across the deletion region using Herculase II (Agilent). This screen used a forward primer upstream of the first CRISPR-Cas9 target site and a reverse primer downstream of the second CRISPR-Cas9 target site (5F and 3R, respectively). Primer information can be found in ***Supplementary Methods***. A PCR product would indicate that at least one allele was deleted from the clone, as the primers in the unedited genome would be too far apart to produce a product of the expected size (***Supplementary Figure 1A, B***). Proceeding with the clones showing at least one allele deleted, a second reaction was run using primers found inside the region to be deleted. These primers are identified as intF and intR. Clones missing this band showed that both alleles had the region deleted, and the segment of DNA was gone (***Supplementary Figure 1A, B***). Homozygous deletion clones were then transferred to Matrigel (Corning, Inc.) and cultured with mTESR Plus (STEMCELL Technologies) for expansion and frozen in vials using Bambanker freezing media.

We used 30× whole-genome sequencing (WGS) to confirm the genomes of the selected clones. WGS libraries were prepared from ∼1μg DNA using Illumina TruSeq PCRfree DNA sample preparation kits targeting an insert size of 350 bp. Library preparation was performed according to the manufacturer’s instructions. The protocols were automated using an Agilent NGS workstation and Beckman Coulter Biomek FXp. WGS clustering was done using cBot and paired-end sequencing with 150 bp read length was performed on Illumina NovaSeq6000 S4. Raw WGS reads were aligned to the GRCh37 human reference genome using BWA-MEM (Li and Durbin 2009, 2010; Li 2013). Variant calls were generated using the GATK pipeline (McKenna et al. 2010). Deletions were visualized in IGV (Robinson et al. 2011; Thorvaldsdóttir et al. 2013; Robinson et al. 2017, 2023).

### Differentiation of iPSCs into multiple schizophrenia-relevant cell types

We first differentiated iPSCs into neural precursor cells (NPCs) and further differentiated NPCs into medium spiny neurons (MSNs) or excitatory glutamatergic neurons (EGNs; ***Supplementary Methods***) (Jacobson 1985; Tam and Keung 2022; Espuny-Camacho et al. 2013). We generated three technical replicates (n = 3; 1-5 million cells per replicate) for each cell type. Following differentiation, we genotyped all replicates and confirmed the lack of off-target variants (UNC HTSF).

### Hi-C Library Preparation

We used the Arima Hi-C Kit to generate genome-wide Hi-C libraries and assess differences in chromatin organization between the two selected deletions and wild-type isogenic controls. We followed the Arima-HiC Kit for Mammalian Cell Lines user guide for cryopreserved cells (Material Part Number: A510008; Document Part Number: A160134 v01). Cryopreserved cell pellets were thawed in a pre-warmed water bath set to 37°C for 90 seconds. Cell pellets were then thawed and crosslinked with 2% formaldehyde in PBS (Sigma-Aldrich, MA), samples were aliquoted and stored at –80°C. Sample DNA concentration was quantified via the Qubit dsDNA High Sensitivity assay (Invitrogen) before proceeding with library preparation using the Arima Library Prep Module (Material Part Numbers: A510008, A303011; Document Part Number: A160432 v01), which outlines the DNA fragmentation and size selection, biotin enrichment, library preparation of enriched ligation products, and final library amplification with purification beads. Briefly, we fragmented DNA using the Covaris S220 focused ultrasonicator for 70 seconds andDNA fragments were then analyzed through a dsDNA Assay 2100 Bioanalyzer (Agilent) to confirm appropriate fragment size distribution centered around 400 bp. Afterwards, we used CleanNGS Beads (Bulldog Bio, NH) to perform DNA Size Selection at a ratio of 0.85-0.56 which enriched for fragments between 200-600 bp (Bulldog CleanNGS SPRI size selection protocol v3). We then assessed sample concentration using dsDNA High Sensitivity Qubit assay. Final library amplification included sample specific indexes (Arima Genomics and xGenIDT, Integrated DNA Technologies, IA). All samples we subjected to five amplification cycles (C1000 Touch Thermal Cycler, BioRad, CA) based on Arima’s DNA input to Cycle Number guidelines. We used AMPure XP Beads (Beckman Coulter, CA) to purify the amplified libraries and assessed their concentrations and fragment size distribution with dsDNA HS Qubit and an Agilent 2100 Bioanalyzer (500 bp on average), respectively. We used Illumina’s NovaSeq 6000 S4 PE (2×150 bp) to generate a minimum of 833 million paired end reads per sample.

### RNA sequencing

For downstream RNA sequencing work, we first removed media, before lysing cells with 350 µl of Buffer RLT with beta-mercaptoethanol (RNeasy Mini kit, Qiagen), resuspending these, and transferring lysate to a 1.5 mL tube, before storing them at –80°C. We extracted RNA according to the RNeasy Mini kit protocol (Qiagen), followed by in-column DNase treatment. Isolated RNA was eluted using nuclease-free water and stored at -80°C until library preparation. RNA libraries were prepared using the KAPA mRNA stranded RNA-seq kit (Roche). Libraries were checked for quality (RNA integrity >9) and quantified using the 4200 TapeStation (Agilent). We used Illumina’s NextSeq 2000 P3 (PE 2×100bp) to generate a minimum of 122 million paired end reads per sample.

### Bioinformatics

#### Hi-C Data Analysis Pipeline

Raw sequencing reads were assessed for quality using FASTQC (version 0.11.7) (https://www.bioinformatics.babraham.ac.uk/projects/fastqc/) and MultiQC (version 1.28). We implemented the Juicer 2.0 pipeline to analyze the Hi-C sequencing data (Durand et al. 2016b). Reads were mapped to the human reference genome version GRCh38/hg38. The pipeline output consisted of .hic files containing contact matrices at multiple resolutions (2.5 Mb to 5 kb). These .hic files included both raw contact counts and matrices normalized using the Vanilla Coverage (VC) method. Juicer automatically merged replicates for each condition (NPC, MSN, ENG, for WT and DEL conditions), facilitating consistent and reproducible downstream analyses. Quality control ensured that more than 80% of reads aligned successfully, with chimeric and ambiguous reads comprising less than 20% of total reads and unmapped reads remaining below 6%. The resulting .hic files were visualized using Juicebox (Durand et al. 2016a) and were subsequently used for downstream analyses, including higher-order chromatin structure assessments such as A/B compartment identification, TAD, subTAD, and TAD-b annotation, and chromatin loop calling using additional tools from the Juicer suite.

### Hi-C Reproducibility Assessment Using HiCRep

We used HiCRep (v3.3.0) to assess the reproducibility of Hi-C contact maps and quantify structural differences across conditions (Lin et al. 2017). HiCRep uses the Stratum-Adjusted Correlation Coefficient (SCC) to provide a more sensitive and accurate comparison of contact matrices (Lin et al. 2017). For comparison, we also performed Pearson correlation analysis as a secondary approach to evaluate consistency between contact maps.

After converting .hic files into .cool files using hicstraw (Durand et al. 2016a) and the Cooler toolkit (Chen et al. 2019), we used HiCRep to compute SCC scores between all pairwise comparisons of conditions across different cell types. A smoothing window size of 1 was applied, and the analysis was restricted to contacts within a 500 kb genomic distance to avoid artifacts due to matrix sparsity. Instead of downsampling, each matrix was normalized by its total contact count to account for differences in sequencing depth. SCC scores were calculated for each chromosome, and the median SCC value across chromosomes was computed per sample to obtain a robust summary statistic. This approach provided a quantitative measure of reproducibility and 3D chromatin structural similarity, enabling more accurate comparisons between cell types. The analysis validated the quality of the Hi-C data, ensured technical consistency across replicates, and captured biological variation associated with cell type differences and differentiation.

### Identification of TADs and TAD-bs

We employed two complementary approaches to identify topologically associating domains (TADs) and their boundaries (TAD-bs): the Arrowhead algorithm from the Juicer Tools suite (Rao et al. 2014) and the insulation score method implemented in FAN-C (version 0.9.28) (Kruse et al. 2020). We ran Arrowhead (Juicer Tools v2.20.00) on .hic files generated by Juicer and normalized using the VC method. Analyses were conducted at 50 kb resolution, which provides a good trade-off between resolution and matrix sparsity. To complement Arrowhead, we also used the insulation score method implemented in FAN-C (Kruse et al. 2020), which identifies TAD-bs based on local decreases in interaction frequency. We used Hi-C data binned at 50 kb resolution and applied a 500 kb sliding window. For each bin, the total contact frequency within the 500 kb × 500 kb square centered on the diagonal was calculated. Bins near the ends of chromosomes without sufficient flanking regions were excluded from the analysis. The insulation scores were computed genome-wide using .hic files as input. Local minima in the insulation score profile, referred to as insulation valleys, were considered candidate TAD boundaries.

### Compartment A/B Identification

To explore large-scale chromatin organization, we performed A/B compartment analysis using the FAN-C toolkit (version 0.9.28) (Kruse et al. 2020). Chromatin compartments represent genome-wide spatial partitioning into transcriptionally active (A) and inactive (B) domains and are typically observed as a plaid-like pattern in Hi-C contact matrices (Lieberman-Aiden et al. 2009). FAN-C implements a series of matrix transformations and eigenvector-based analyses to identify and quantify these compartments (Kruse et al. 2020). We provided genomic GC content information for GRCh38/hg38, such that positive values correspond to GC-rich, gene-dense A compartments. This ensured consistent and biologically meaningful compartment labeling across all chromosomes.

We also generated correlation matrices to enable quality control and reproducibility analyses. For our study, Hi-C data binned at 100 kb and 1 Mb resolution were used as input, derived from .hic files.

### Compartment Assignment and Flip Correction Using RNA-seq Data

Although most chromosomes displayed clear A/B compartment structure based on the output from FAN-C (Kruse et al. 2020), some chromosomes were difficult to interpret in terms of the directionality of the first eigenvector. To biologically validate and orient the A/B compartment signs, we used matched RNA-seq data as an independent indicator of transcriptional activity. The A compartment is typically associated with euchromatin, open chromatin states, high gene density, and active transcription, while the B compartment corresponds to heterochromatin and transcriptionally inactive regions. Therefore, regions with high gene expression were expected to align with the A compartment, providing a rationale for confirming or correcting the eigenvector orientation.

To compare RNA expression with Hi-C–derived compartments, expression values were binned to match the 1 Mb resolution used in the compartment analysis. Using bedtools (Quinlan and Hall 2010), we averaged the expression levels of all genes within each 1 Mb non-overlapping genomic bin. This approach allowed us to directly compare transcriptional activity with the A/B compartment values derived from FAN-C and validate compartment assignments across the genome.

### Chromatin Loop Detection and Differential Analysis

Chromatin loop detection was performed using the HiCCUPS algorithm (version 1.9.9) (Rao et al. 2014) within Juicer (Durand et al. 2016b). Hi-C contact matrices were generated from .hic files using Juicer with Vanilla Coverage (VC) normalization to adjust for sequencing depth and technical biases. Loop detection was conducted at 5 kb, 10 kb, and 25 kb resolutions to capture interactions across different genomic scales. For each sample, peak detection was carried out individually. We only considered interactions spanning 35 kb to 2 Mb, where the lower limit avoids detection of self-ligation events. HiCCUPS applies a modified Benjamini-Hochberg correction to control the false discovery rate (FDR) within each chunk. A minimum of 16 reads in the lower-left neighborhood was required to estimate background; otherwise, the window was incrementally expanded to a maximum width of 20 to ensure robustness in sparse regions. Detected loops within 20 kb of each other were clustered into a single consolidated call to reduce redundancy and improve confidence.

### Hi-C Loop Annotation and Classification

HiCCUPS loop calls generated by Juicer v2.0 were annotated to integrate regulatory element information and to exclude regions overlapping known genomic artifacts. Regulatory element annotations were obtained from the ENCODE registry of candidate cis-regulatory elements (cCRE Registry version 3, GRCh38/hg38) (ENCODE Project Consortium et al. 2020). Promoter-like signatures (PLS) were classified as promoters, while proximal and distal enhancer-like signatures (pELS and dELS) were classified as enhancers. Anchors overlapping ENCODE blacklist regions were removed. Anchors were annotated according to their overlap with cCREs and loop type was inferred from the combination of both anchors, yielding promoter–enhancer, promoter–promoter, enhancer–enhancer, or other categories. A dynamic Bonferroni correction was applied to control for multiple testing, with the adjusted significance threshold calculated as α/N (α = 0.05, N = total number of detected loops). Loops with fdrBL values below this threshold were considered high-confidence interactions.

### Differential Analysis of Chromatin Organization

#### A/B Compartment Comparison Across Conditions

We compared A/B chromatin compartments across conditions using two complementary approaches. First, we identified polarity switches (A-to-B or B-to-A) at 1 Mb resolution. Second, we applied the statistical framework provided by dhHiC (Chakraborty et al. 2022) at 100 kb resolution. For dhHiC-based analysis, we preprocessed the Hi-C data in .cool format. Contact matrices and bin definitions were extracted using the cooler toolkit (Open2C et al. 2024). Only for the dhHiC-based differential compartment analysis, bins corresponding to non-standard chromosomes (chrM, chrX, chrY) were excluded to minimize technical artifacts, we excluded bins from non-standard chromosomes generating a list of their bin IDs and removing interactions involving these bins, retaining only autosomal interactions. We then computed the first principal component (PC1) values using dhHiCcall to assign compartment identities. dhHiC performed pairwise comparisons between samples to detect regions with significant compartment shifts.

#### Differential TAD analysis across conditions

To detect differences in TAD organization between conditions, we used DiffDomain (Hua et al. 2024), a statistical method specifically developed to identify significant changes in TAD-bs from Hi-C data. For this analysis, we used TAD-b files previously identified in each condition using Arrowhead (Durand et al. 2016b) and conducted differential analysis at 50 kb. We used as input the TAD boundaries from one condition along with the corresponding Hi-C files from both conditions. We applied the default p-value cutoff of 0.05 to identify significantly altered TAD regions between conditions. The resulting output highlighted genomic loci where TAD organization was significantly different, providing insights into how 3D genome architecture is altered in response to specific biological states.

#### Differential chromatin looping analysis across conditions

We used the hiccupsdiff (Durand et al. 2016b) function from the Juicer Tools suite to identify differential chromatin loops between experimental conditions at a 5 kb resolution. After loops were independently detected for each sample using HiCCUPS, hiccupsdiff was used to systematically compare each experimental group against its corresponding control group. This comparison enabled the detection of statistically significant changes in chromatin loop structures, identifying interactions that were either gained or lost between conditions.

### RNA Sequencing Quantification and Differential Expression Analysis

RNA sequencing (RNA-seq) data were processed using a transcript-level quantification approach followed by gene-level differential expression analysis. Transcript abundance was quantified using Salmon (Patro et al. 2017), a lightweight, alignment-free method that enables rapid and accurate transcript quantification. For this analysis, a custom Salmon index was built using the GENCODE v21 reference transcriptome, limited to protein-coding transcripts only. We retained transcript-level quantification (txOut = TRUE) to enable the analysis of protein-coding transcripts with higher resolution and flexibility in downstream modeling.

To complement and validate the quantification results, an alternative pipeline based on traditional alignment was also implemented using the STAR aligner (Dobin et al. 2013), allowing for direct comparison between alignment-free and alignment-based RNA-seq workflows. This dual approach ensured consistency and robustness in transcript quantification prior to downstream analysis.

Principal component analysis (PCA) was performed using normalized transcript counts from all samples to assess global sample clustering and identify potential outliers. Following this initial quality assessment, differential expression analysis was conducted by comparing only the samples carrying genomic deletions with their corresponding matched controls, stratified by cell type. This comparison strategy allowed for the identification of gene expression changes specifically associated with the two genomic deletions, minimizing confounding effects from unrelated biological or technical variation. Differential expression was carried out using DESeq2 (Love et al. 2014), which models count data using negative binomial distributions and adjusts for differences in sequencing depth. Genes with an adjusted p-value (FDR) below 0.05 were considered significantly differentially expressed.

### Integration of chromatin looping and gene expression

Differential chromatin loops identified by HICCUPSDiff were assigned to nearby protein-coding genes based on their genomic coordinates (hg38). A gene was considered associated with a loop when either loop anchor overlapped a genomic window extending ±5 kb from the annotated gene coordinates, corresponding to the 5-kb Hi-C bin size used for loop detection. Genes associated with either anchor were retained. Each loop was subsequently annotated as promoter-enhancer, enhancer-enhancer, or other regulatory interaction. To identify genes with potential functional consequences of chromatin remodeling, loop-associated genes were intersected with differentially expressed genes (DEGs; Benjamini-Hochberg adjusted P < 0.05) identified by RNA sequencing. The resulting overlap defined the set of loop-associated DEGs used for downstream analyses.

### Pathway Enrichment Analysis

Pathway enrichment was performed using an in-house computational workflow designed to assess whether specific biological pathways were over-represented among genes identified as differentially expressed or associated with altered chromatin interactions in each comparison. The foreground gene set for each analysis consisted of unique Ensembl gene identifiers derived from the corresponding experimental condition.

The background gene set was defined from the expressed protein-coding genes detected in control samples (expression >1 in both control replicates). Transcripts per million (TPM) values from two control samples (n = 2) were extracted, and the maximum TPM was calculated for each gene. Genes with maxTPM ≥ 1 were retained as expressed; a threshold commonly used to capture reliably transcribed genes while minimizing noise from low-abundance transcripts. Prior to analysis, duplicate gene entries were removed, and the list was further restricted to protein-coding genes based on Ensembl gene-biotype annotation.

Functional gene sets were obtained from a consolidated reference database, which integrates multiple curated annotation resources (***Supplementary Table 4***). These include SynGO Biological Process and Cellular Component annotations (Koopmans et al. 2019), Gene Ontology Biological Process, Cellular Component, and Molecular Function terms (The Gene Ontology Consortium 2019), RNA-binding protein target lists (CELF4 and RBFOX1–3), disease-associated gene sets, drug-target gene sets, evolutionary constraint categories (Sullivan et al. 2023), general gene-annotation resources, WikiPathways, and expression-based gene groups. Only these predefined categories were retained for enrichment testing to ensure systematic and comparable analyses across datasets.

Enrichment was evaluated using a hypergeometric over-representation test, which measures whether the overlap between the foreground gene list and each pathway exceeds that expected by chance. To avoid unstable estimates, gene sets with fewer than five overlapping foreground genes were excluded. Multiple-testing correction was performed using the Benjamini–Hochberg false discovery rate (FDR) procedure, and pathways with FDR < 0.01 were considered significantly enriched. For each significant pathway, the full list of overlapping foreground genes was extracted to support downstream biological interpretation, clustering analyses, and visualization.

### Data availability

These data have been deposited in the National Center for Biotechnology Information Gene Expression Omnibus, and the accession number is pending.

### Code Availability

https://github.com/giustirodriguezlab/3D-genome-impacts-of-TAD-boundary-disrupting-ultra-rare-structural-variants-identified-in-scz

## Results

Our study successfully modeled and characterized two schizophrenia-associated TAD-boundary-disrupting deletion SVs (SV Del10: hg38 chr10:44,826,494–44,834,460; SV Del13: hg38 chr13:51,088,404–51,094,873) identified in our schizophrenia whole-genome sequencing study (Halvorsen et al. 2020) (***Figure 1A***). These variants were selected based on multiple lines of genetic and functional evidence, including predicted disruption of TAD-bs, evolutionary conservation, regulatory annotations, and association with schizophrenia risk loci (see ***Methods***).

**Figure 1.**
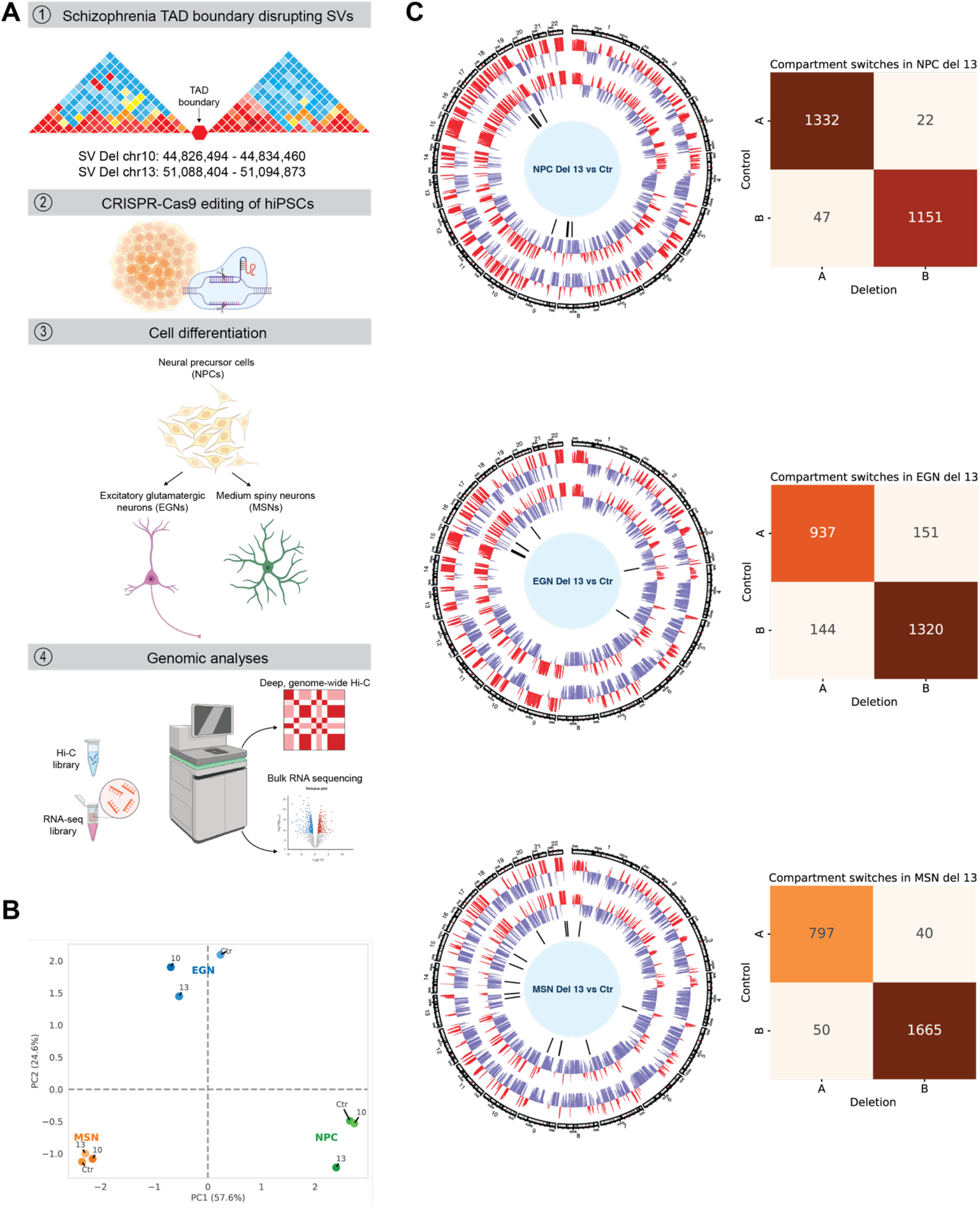
Experimental design and global characterization of 3D genome organization. (**A**) Schematic of our experimental design. Two schizophrenia-associated TAD boundary-disrupting deletions (SV Del10 and SV Del13; hg38 coordinates shown) were engineered into an isogenic human iPSC line using CRISPR-Cas9. Cells were differentiated into neural progenitor cells (NPCs), excitatory glutamatergic neurons (EGNs), and medium spiny neurons (MSNs) and profiled by in situ Hi-C and bulk RNA sequencing. (**B**) Principal component analysis (PCA) of A/B compartment eigenvectors extracted using FAN-C at 1-Mb resolution. Samples are colored by cell type (NPC, green; EGN, blue; MSN, orange) and labeled by genotype (CTR, control; 10, SV Del10; 13, SV Del13). Dashed lines indicate the origin (PC1 = 0, PC2 = 0). (**C**) Genome-wide A/B compartment analysis at 1-Mb resolution in control and SV Del13 samples across NPCs, EGNs, and MSNs. Circos plots show chromosome ideograms, followed by A/B compartment profiles for the SV Del13 sample and its matched control, and an inner track highlighting genomic regions undergoing A↔B compartment switching. The accompanying color-coded contingency table summarizes the distribution of compartment transition categories (A to A, A to B, B to A, and B to B) for each cell type, with darker shading indicating a higher proportion of genomic regions in each transition category.

We sought to determine how disruption of two distinct TAD-bs influences chromatin architecture and gene regulation during neuronal differentiation. To address this, we generated isogenic CRISPR-Cas9-engineered human iPSC lines carrying each deletion (***Figure 1A***). Three independent clones were established for each deletion, and 30× whole-genome sequencing confirmed genome integrity and the absence of detectable off-target events (***Supplementary Figure 1C, D***). The engineered lines were differentiated into neural progenitor cells (NPCs), excitatory glutamatergic neurons (EGNs), and medium spiny neurons (MSNs), and profiled using high-resolution in situ Hi-C and bulk RNA sequencing to characterize the effects of each deletion on 3D genome organization and transcription across neuronal lineages (***Figure 1A***).

### Cell-type identity is the primary determinant of 3D genome organization across SV deletion backgrounds

To establish a robust foundation for comparative analyses of chromatin architecture across cell types and SV deletion backgrounds, we first assessed Hi-C library quality. Juicer quality-control metrics indicated consistently high sequencing and mapping performance across all samples, supporting reliable downstream analyses (***Supplementary Table 1***). Across all samples, more than 87%–91% of reads were successfully aligned as paired reads, exceeding the >80% threshold expected for high-quality Hi-C libraries. The proportion of reads without detected chimeric junctions was consistently low (0.32%–0.48%), and unmapped reads accounted for less than 6% of total reads in all samples, meeting recommended quality benchmarks. Hi-C contacts comprised 63.6%–66.7% of total reads and represented 69.9%–74.2% of all alignable reads, exceeding the expected >60% threshold for productive Hi-C libraries. Moreover, long-range cis interactions (>20 kb), a key indicator of informative chromatin contacts, accounted for approximately 34%–42% of unique Hi-C contacts, meeting or exceeding the >40% benchmark for datasets warranting deep sequencing. Inter- and intra-chromosomal contact proportions were consistent across genotypes and cell types, indicating uniform library complexity and digestion–ligation efficiency. Collectively, these metrics confirm that all Hi-C libraries met or exceeded recommended standards and were suitable for high-resolution analyses of chromatin compartments, TADs, and looping interactions.

In addition to standard Juicer quality-control metrics, we evaluated reproducibility across biological replicates and cell types and observed exceptionally high concordance (***Supplementary Figure 2A***). Stratum-adjusted correlation coefficient (SCC) values showed near-perfect reproducibility within the same cell type, with correlations approaching or exceeding 0.99 across both control and SV-deletion samples, demonstrating strong consistency across genotypes (***Supplementary Figure 2A***). Correlations between different cell types were slightly lower (approximately 0.90–0.96), reflecting pronounced cell-type–specific chromatin organization.

Importantly, SV Del10 and SV Del13 samples clustered more closely with their corresponding cell-type controls than with other lineages, indicating that cell-type identity, rather than SV status, is the dominant source of variation in 3D genome architecture. Moreover, EGN and MSN samples exhibited higher mutual correlations compared to NPCs, suggesting that differentiated neuronal lineages share more similar 3D genome architectures than progenitor states (***Supplementary Figure 2A***). Together, these results confirm the robustness of the Hi-C datasets and demonstrate strong preservation of cell-type–specific chromatin structure across structural variant backgrounds. No systematic differences in QC metrics were observed between SV-deletion and control samples, indicating that downstream differences reflect biological rather than technical variation.

### Global compartment architecture remains largely preserved following SV deletions

With high-quality and reproducible Hi-C datasets established, and after observing that samples clustered primarily by cell type rather than genotype, we next investigated whether SVs altered higher-order chromatin architecture within each neuronal lineage. Principal component analysis of both compartment eigenvectors (***Figure 1B***) and Hi-C contact matrices (***Supplementary Figure 2B***), demonstrated that cell type was the primary determinant of genome-wide chromatin organization, whereas SV-associated differences contributed a secondary source of variation. Therefore, all downstream analyses compared each SV-deletion sample with its matched control within the corresponding neuronal lineage.

We first examined A/B compartment organization, which partitions the genome into large-scale active (A) and inactive (B) chromatin domains associated with transcriptional activity, at both 1-Mb and 100-kb resolution. At 1-Mb resolution, both SV Del10 and SV Del13 produced only modest alterations in compartment organization, indicating that global chromatin compartmentalization remained largely preserved following either deletion (***Figure 1C***). Increasing the resolution to 100 kb revealed a greater number of compartment transitions, suggesting that the effects of the deletions were primarily localized rather than reflecting widespread genome-wide reorganization (***Supplementary Figure 2C***).

Among the three neuronal lineages, EGN neurons exhibited the strongest compartmental response, with approximately 11.6% of genomic bins switching compartments following SV Del13 (***Figure 1C***) and 14.6% following SV Del10, compared with approximately 3% in MSN cells and 2% in NPCs under either deletion (***Supplementary Figure 3A-C***). The directionality of compartment transitions also varied according to both cell type and SV. In MSN cells, SV Del13 induced nearly equal proportions of A-to-B and B-to-A transitions (∼1.4% each), whereas SV Del10 was characterized predominantly by B-to-A switching. EGN neurons displayed relatively balanced transitions in both directions following either deletion. In contrast, NPCs exhibited predominantly B-to-A transitions under both SV Del10 and SV Del13. Together, these findings demonstrate that although both SV deletions largely preserve global compartment architecture, they induce modest, lineage-specific remodeling of chromatin compartments, with differentiated excitatory neurons exhibiting the greatest susceptibility to compartment reorganization.

### SV deletions weaken local insulation and alter boundary positioning without disrupting TAD structure

To determine whether the SV affected chromatin organization at a finer scale, we next examined topologically associating domains (TADs) and boundary insulation, which are fundamental units of 3D genome organization that constrain enhancer-promoter interactions and help maintain proper gene regulation. TADs were identified using Arrowhead at 50-kb resolution, and insulation scores were calculated using FAN-C to quantify changes in boundary strength.

Despite the presence of the deletions, Hi-C maps showed that domain-scale chromatin organization remained largely unchanged across all three neuronal lineages, with no apparent TAD collapse or fusion at either locus (***Figure 2A-B***). At the SV Del10 locus, the deleted interval lies adjacent to a TAD-b, whereas SV Del13 directly overlaps a boundary (***Figure 2A-B***; ***Supplementary Figure 4A-D***). Across all three neuronal lineages, the overall TAD architecture surrounding both deletions remained intact. However, SV Del10 produced a modest positional shift of the neighboring boundary in MSN and EGN cells, whereas no comparable shift was observed in NPCs, suggesting that deletion of sequence adjacent to a boundary can subtly alter boundary positioning without disrupting domain structure.

**Figure 2.**
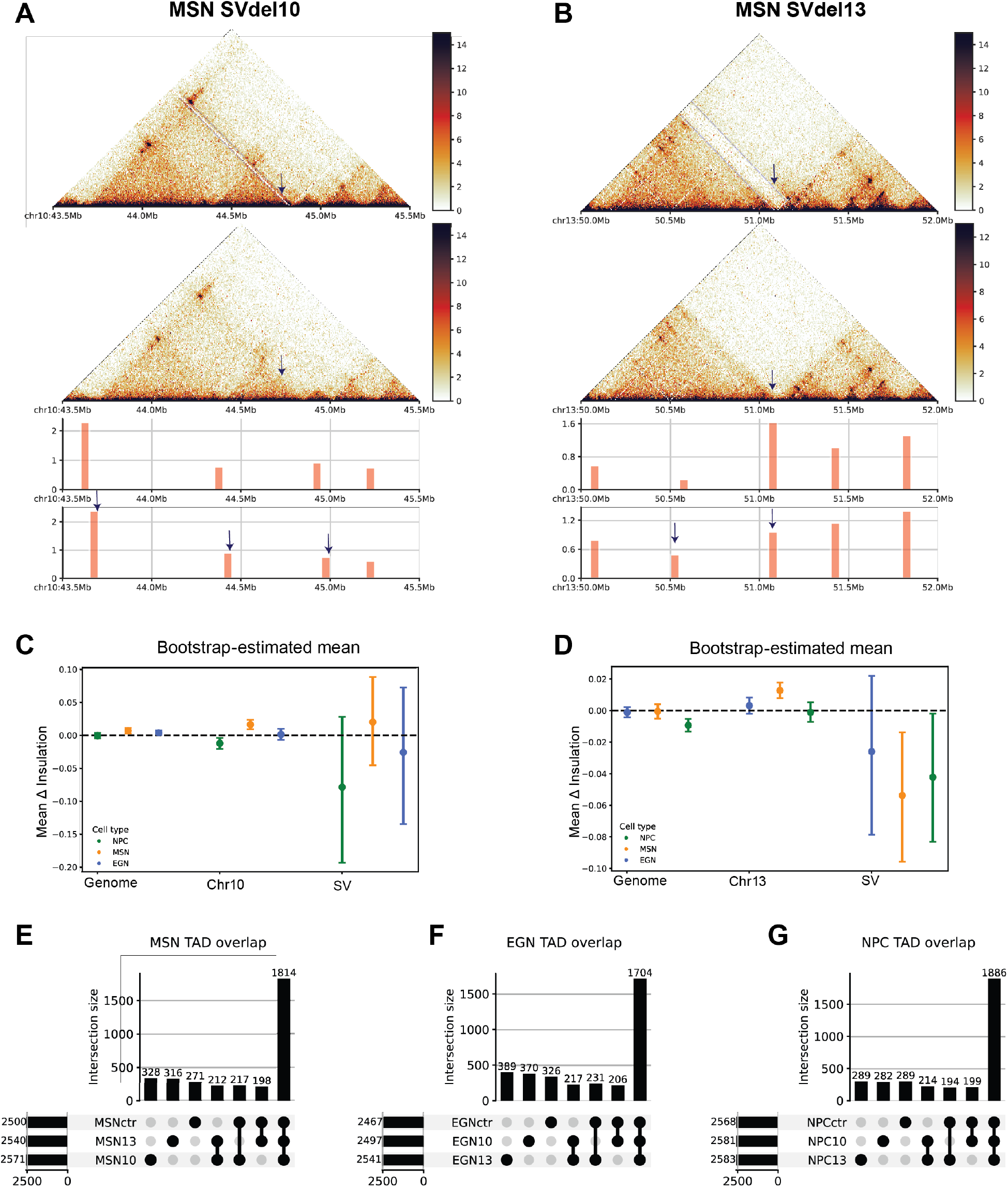
Chromatin insulation and TADs changes induced by SV deletions. (**A–B**) Representative Hi-C contact maps centered on the edited locus in MSN cells for SV Del10 (**A**) and SV Del13 (**B**). The upper panels show the deletion samples, in which the edited region is visible as a gap in contact signal (highlighted by arrows), whereas the lower panels show the corresponding contact maps from the matched control samples. Beneath the contact maps, bar plots of insulation scores are shown separately for the SV deletion (upper panel) and the matched control (lower panel), highlighting visible changes in chromatin insulation surrounding the deleted TAD boundary. (**C–D**) Bootstrap estimates of the mean change in insulation score comparing SV Del10 (**C**) and SV Del13 (**D**) with their corresponding matched controls. Changes were quantified at four genomic scales: genome-wide, chromosome-wide, the chromosome containing the deletion, and the deleted TAD boundary region (SV). Colors denote cell types: NPCs (green), EGNs (blue), and MSNs (orange). (**E–G**) UpSet plots showing the overlap of TAD boundary positions identified in MSNs (E), EGNs (F), and NPCs (G) across control (CTR), SV Del10 (10), and SV Del13 (13) samples. TAD boundaries were considered overlapping when detected within a 50-kb genomic window, corresponding to the resolution used for TAD boundary identification.

To quantify these observations, we analyzed FAN-C differential insulation scores across the genome, the affected chromosome, and the deleted interval (***Figure 2C-D***). For SV Del10, differential insulation remained centered near zero across the deleted interval in all three cell types, with bootstrap confidence intervals overlapping zero and no significant difference relative to the genomic background (Mann-Whitney test, all P > 0.55) (***Figure 2C***). Thus, although SV Del10 altered the apparent position of nearby boundaries in differentiated neurons, it did not measurably weaken boundary insulation.

In contrast, SV Del13 produced a localized reduction in boundary insulation. Genome-wide mean differential insulation remained close to zero in all three neuronal lineages (mean Δ = −0.011 to 0.0004), indicating that boundary insulation was largely preserved at the chromosomal and genomic scales. However, within the deleted interval, EGN and MSN cells exhibited substantially more negative differential insulation (mean Δ = −0.087 and −0.094, respectively), with bootstrap confidence intervals excluding zero and Wilcoxon tests indicating significant local weakening (EGN: p = 0.016; MSN: p = 0.020) (***Figure 2D***). Moreover, insulation changes at the deleted interval were significantly greater than those observed across the remainder of the genome (Mann-Whitney P = 0.010 and 0.007, respectively). NPCs showed a similar trend (mean Δ = −0.054), although neither the confidence interval nor the comparison with the genomic background reached significance.

Finally, we asked whether these local boundary perturbations resulted in reorganization of TADs by comparing domain coordinates between control and deletion samples using positional overlap (***Figure 2E-G***). TAD organization remained highly conserved across all neuronal lineages. NPC cells retained 1,886 shared TADs, with only 282 and 289 domains unique to the SV Del10 and SV Del13 comparisons, respectively. Similarly, MSN cells shared 1,814 TADs, with 217 SV Del10-specific and 198 SV Del13-specific domains, whereas EGN cells retained 1,704 shared TADs compared with 206 and 231 deletion-specific domains.

Overall, more than 85–90% of TADs were conserved across genotypes in every lineage, demonstrating that neither structural variant substantially reorganized domain-scale chromatin architecture. Together, these results indicate that the genomic context of the deletion determines its structural consequence. SV Del13 directly affect a TAD-b, leading to localized weakening of boundary insulation without altering overall TAD organization, whereas SV Del10 lies adjacent to a boundary and primarily causes subtle positional shifts while preserving both boundary strength and domain architecture.

### Localized regulatory loop rewiring represents the primary 3D genome consequence of SV deletions

Although both deletions produced only modest changes in chromatin compartments and largely preserved TAD organization, alterations in boundary positioning and insulation suggested that regulatory interactions might be remodeled at a finer scale. We therefore investigated chromatin looping, the highest-resolution feature of 3D genome organization, to determine whether these local structural perturbations altered enhancer-promoter interactions.

Across all cell types, we detected extensive chromatin looping interactions, of which a subset represented high-confidence regulatory loops linking promoters and distal regulatory elements (***Supplementary Table 1***). In total, approximately 18,000, 20,000, and 13,000 chromatin loops were identified in MSN, NPC, and EGN cells, respectively. Among these, roughly 10,000 loops in MSNs, 9,000 in NPCs, and 8,000 in EGNs overlapped candidate cis-regulatory elements (cCREs), including enhancer-enhancer, promoter-promoter and enhancer-promoter interactions (***Figure 3A-C***). After applying these regulatory annotations, MSN cells contained the largest number of high-confidence regulatory loops (∼5,000), followed by NPCs (∼4,600) and EGNs (∼3,500) (***Supplementary Table 2***). Comparison of loop coordinates between control and SV-deletion samples revealed extensive genotype-specific loop changes, indicating that many loops were detected in only one genotype. However, these positional differences did not necessarily correspond to changes in interaction strength.

**Figure 3.**
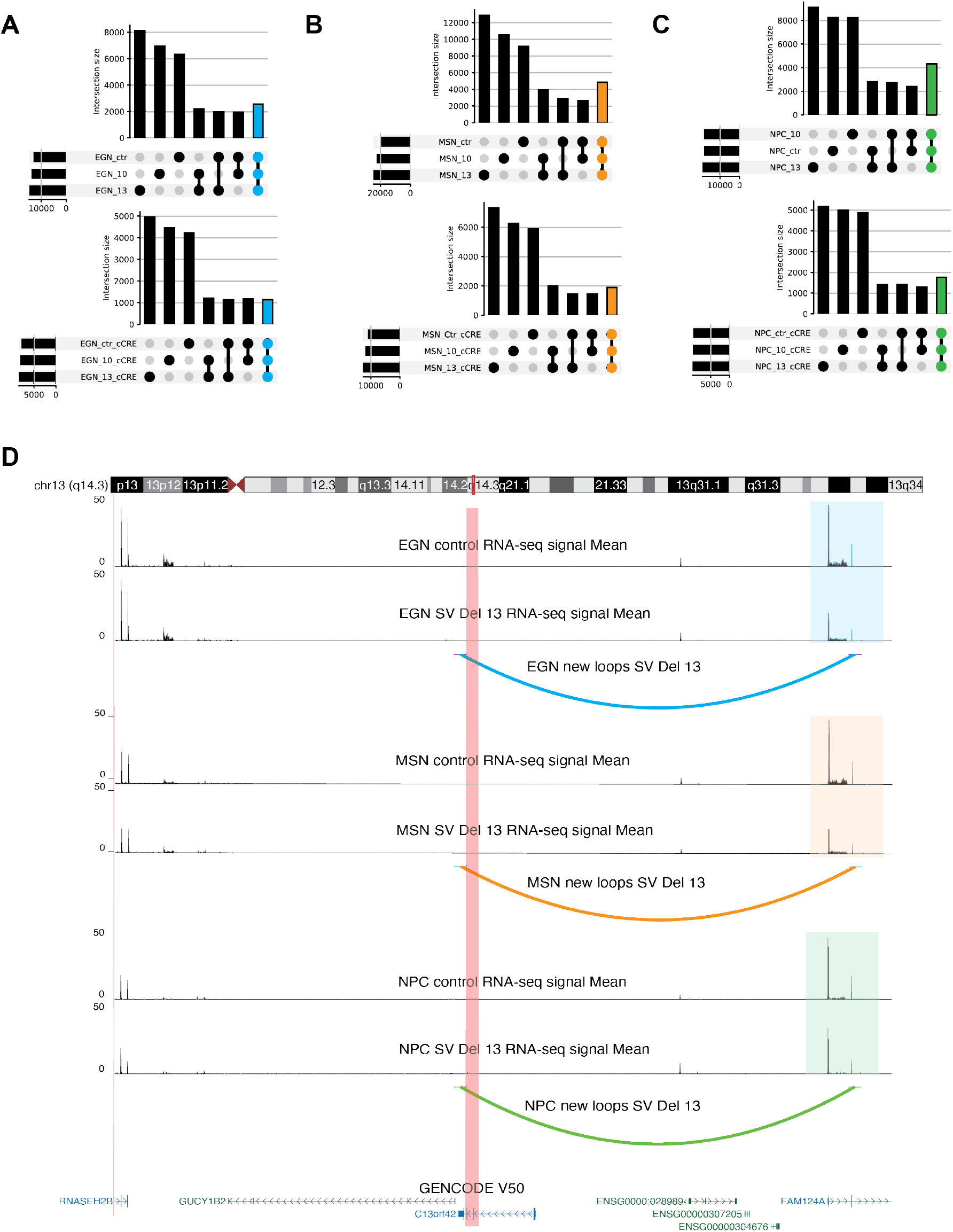
Structural variant-induced chromatin loop remodeling. (**A–C**) UpSet plots showing the overlap of chromatin loops across control (CTR), SV Del10 (10), and SV Del13 (13) samples in EGNs (**A**), MSNs (**B**), and NPCs (**C**). The upper UpSet plot summarizes all chromatin loops detected across the three genotypes, whereas the lower UpSet plot includes only loops classified as candidate cis-regulatory element (cCRE)-associated interactions. Bar colors indicate the number of loops shared among CTR, SV Del10, and SV Del13 based on loop position. (**D**) Integrated view of the genomic region surrounding the SV Del13 locus. The deleted region is highlighted in red on the chromosome ideogram and within the local genomic view. Differential chromatin loops and mean RNA-seq expression levels are displayed for NPCs, EGNs, and MSNs to illustrate the relationship between SV-induced chromatin loop remodeling and local transcriptional changes. The bottom track shows GENCODE v50 gene annotations.

To identify robust alterations in chromatin architecture, we next compared loop interaction strengths using HiCCUPSDiff (***Supplementary Table 2***). Between 1,093 and 1,314 differential loops were detected across individual cell type–deletion comparisons, representing approximately one-quarter to one-third of the total regulatory loop repertoire (***Figure 3A-C***). MSN cells exhibited the largest number of differential loops under both deletions (1,182 for SV Del10 and 1,314 for SV Del13; ***Figure 3B***), whereas NPCs and EGNs showed comparable but slightly fewer remodeled interactions (***Figure 3A, C***). Across all comparisons, enhancer-enhancer interactions consistently represented the largest class of differential loops (690–749 loops; 57–61%), whereas promoter-enhancer interactions accounted for only 99–128 loops (8–11%), with the remaining interactions classified as other regulatory contacts (***Supplementary Table 2***). These results indicate that SV-associated chromatin remodeling occurs predominantly through changes in distal enhancer connectivity rather than widespread rewiring of direct promoter interactions.

Although the overall number of differential loops was similar between the two deletions, the direction of loop remodeling differed. In MSN cells, SV Del13 preferentially generated new chromatin interactions, with 613 gained loops compared with substantially fewer lost loops, whereas SV Del10 produced a more balanced distribution of loop gains and losses. A similar gain-biased pattern was observed in EGN cells, where SV Del13 generated 543 gained loops among 869 differential regulatory interactions, while SV Del10 again showed nearly equal numbers of gained and lost loops. In contrast, NPC cells displayed comparable proportions of gained and lost loops under both deletions (824 in SV-del10 and 854 in SV-del13), suggesting that the bias toward the emergence of new loops by SV Del13 is most pronounced in differentiated neuronal lineages (***Supplementary Table 3***).

To determine whether the deletions directly affected local 3D-genome organization, we examined loops anchored within the deletion intervals. SV-del13 consistently generated a novel loop in all three cell types, linking the deleted region to genes that also exhibited altered expression (***Figure 3D***). In contrast, SV-del10 did not produce a consistent looping change at the deletion site. These findings are consistent with the insulation analyses presented above: SV Del13, which weakened boundary insulation, also generated a reproducible novel loop at the disrupted boundary, whereas SV Del10 induced only subtle boundary repositioning without consistent loop remodeling at the deletion locus.

### Integration of 3D genome and transcriptomic data identifies loop-associated gene dysregulation

The identification of SV-induced changes in local chromatin looping, including the formation of a novel loop at the SV Del13 locus that was associated with altered expression of nearby genes (***Figure 3D***), prompted us to investigate the broader transcriptional consequences of chromatin loop remodeling. We integrated differential chromatin loops with RNA-seq data by intersecting genes associated with statistically significant gained or lost loops with differentially expressed genes (DEGs). Consistent with the Hi-C data, principal component analysis (PCA) of the RNA-seq profiles showed that cell-type identity was the dominant source of transcriptional variation (***Figure 4A***).

**Figure 4.**
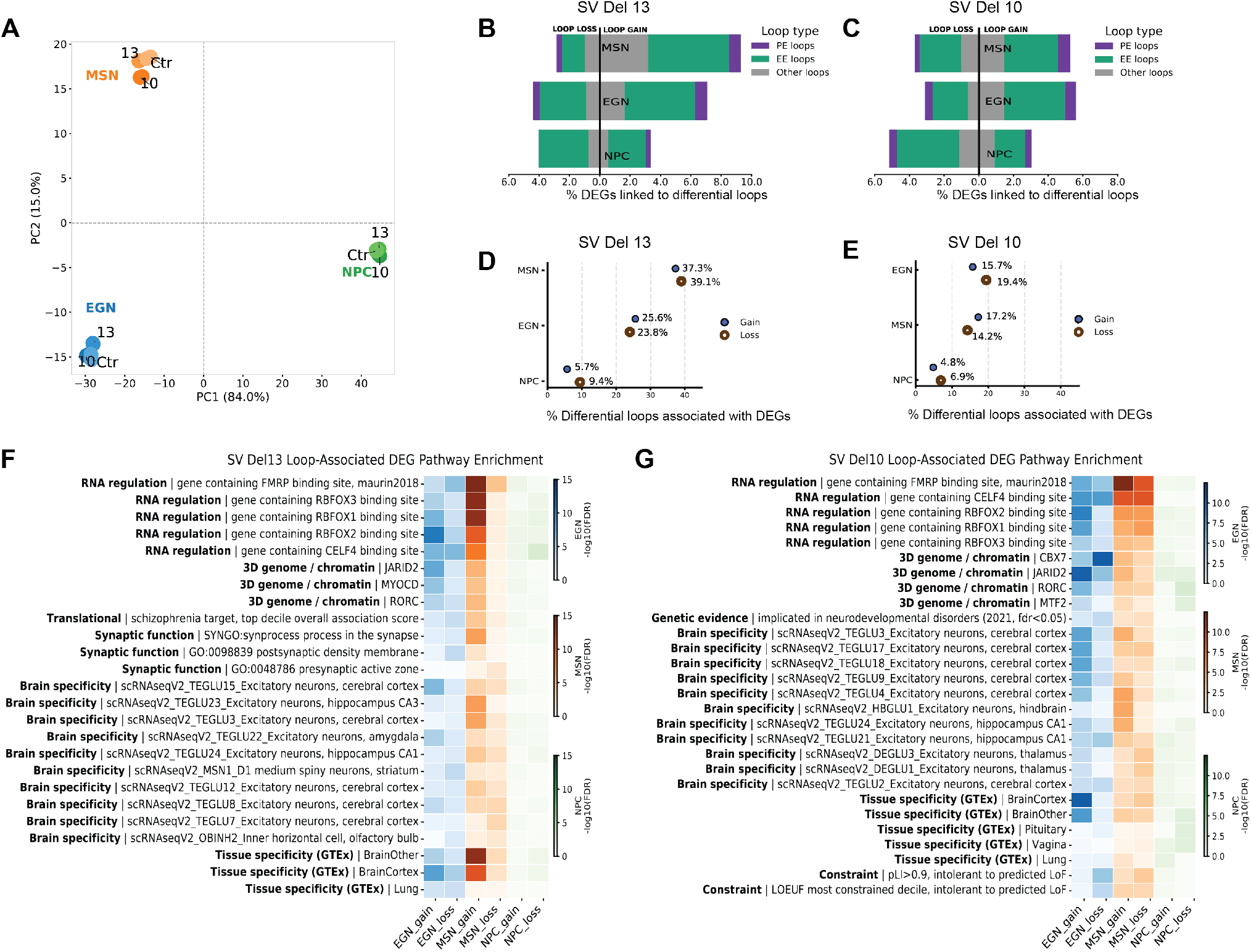
Integration of 3D genome organization and transcriptomics identifies loop-associated gene dysregulation and convergent neuronal regulatory programs. **(A)** Principal component analysis (PCA) of RNA-seq expression profiles across all samples. Samples are colored by cell type (NPC, green; EGN, blue; MSN, orange) and labeled by genotype (Ctr, control; 10, SV Del10; 13, SV Del13). Dashed lines indicate the origin (PC1 = 0, PC2 = 0). **(B–C)** Proportion of differentially expressed genes (DEGs) associated with differential chromatin loops in SV Del13 **(B)** and SV Del10 **(C)** across NPCs, EGNs, and MSNs. Stacked bars classify loop-associated DEGs according to loop type (promoter-enhancer, enhancer-enhancer, and other interactions). Values to the left and right of each bar indicate the percentage of DEGs associated with lost and gained chromatin loops, respectively. **(D–E)** Percentage of differential chromatin loops associated with at least one differentially expressed gene in SV Del13 (D) and SV Del10 (E) across NPCs, EGNs, and MSNs. Separated into gained and lost chromatin loops. (**F–G**) Heatmaps showing the top significantly enriched pathways among loop-associated DEGs for SV Del13 (**F**) and SV Del10 (**G**). Pathway enrichment was assessed using a hypergeometric test with Benjamini–Hochberg correction (FDR < 0.05). Rows represent the top 10 significantly enriched pathways identified across all comparisons, grouped into functional categories (RNA regulation, 3D genome/chromatin, translational, genetic evidence, synaptic function, brain specificity, and tissue specificity). Columns correspond to gained- and lost-loop-associated DEGs in NPCs, EGNs, and MSNs. Color intensity represents enrichment significance (−log10 FDR), with separate color scales for each cell type.

We then quantified the overlap between genes connected to differential chromatin loops and DEGs (***Figure 4B-C***). Across all neuronal lineages, approximately 8–12% of DEGs were connected to differential chromatin loops, indicating that a subset of transcriptional changes occurred within regions undergoing local 3D genome remodeling. The proportion of loop-associated DEGs differed by both cell type and deletion. SV Del13 showed the strongest effect in differentiated neurons, where 11.1% of DEGs in EGN cells and 11.6% in MSN cells were associated with altered loops (***Figure 4B***), compared with 8.4% and 8.6%, respectively, following SV Del10 (***Figure 4C***). In contrast, NPC cells exhibited similar proportions under both deletions (7.5–8.1%) (***Figure 4B-C***).

Reciprocally, only a subset of differential chromatin loops was associated with altered gene expression, indicating that not all structural remodeling resulted in detectable transcriptional consequences. Although SV Del13 showed a predominance of gained loops, particularly in EGN and MSN cells, gained and lost loops associated with DEGs were more evenly distributed. Moreover, a greater proportion of altered loops was associated with DEGs in SV Del13 than in SV Del10 (***Figure 4D-E***). For SV Del13, although the relationship between loop remodeling and transcription was even stronger in differentiated neurons (***Figure 4D***). In EGN cells, 25.6% of gained loops and 23.8% of lost loops were associated with DEGs, increasing to 37.3% and 39.1%, respectively, in MSN cells. By comparison, only 5.7% of gained loops and 9.4% of lost loops in NPCs were linked to differential gene expression (***Figure 4D***), indicating that SV-induced chromatin loop remodeling is substantially more likely to have transcriptional consequences in differentiated neuronal lineages than in neural progenitor cells. A similar trend was observed for SV Del10, 15.7% of gained loops and 19.4% of lost loops in EGN cells were associated with at least one DEG, while the corresponding values in MSN cells were 17.2% and 14.2%, respectively. In NPCs, however, only 4.8% of gained loops and 6.9% of lost loops were linked to DEGs (***Figure 4E***).

We also examined the types of regulatory interactions associated with loop-linked DEGs. Across both deletions and all three cell types, approximately two-thirds (65–66%) of loop-associated DEGs were connected through enhancer-enhancer interactions, whereas only ∼10% involved promoter-enhancer loops (***Supplementary Table 3***), suggesting that structural variant-induced chromatin remodeling primarily affects distal enhancer networks rather than direct promoter contacts.

### SV-induced chromatin loop remodeling converges on common neuronal regulatory programs

The convergence of differential chromatin looping and gene expression changes enabled us to investigate the biological pathways most directly affected by SV-induced chromatin remodeling. We therefore performed pathway enrichment analysis using the identified genes from the previous section (***Figure 4F*, *G***). Despite the distinct genomic mechanisms underlying each SV and the cell type-specific response, loop-associated DEGs converged on a common set of neuronal regulatory programs, indicating that these two patterns of local 3D-genome remodeling ultimately perturb similar biological processes. Consistent with the chromatin looping analyses, these enrichments were driven predominantly by genes associated with gained loops following SV Del13, whereas SV Del10 showed comparable contributions from gained- and lost-loop-associated genes. Across both deletions, the strongest enrichments involved neuronal RNA regulatory networks, including targets of the RNA-binding proteins RBFOX1, RBFOX2, RBFOX3, CELF4, and FMRP, together with chromatin regulatory pathways and brain-specific gene sets derived from GTEx ***(Figure 4F, G***). Overall, these enrichments were substantially stronger in differentiated neurons (EGN and MSN) than in NPCs, indicating that schizophrenia-associated SV-driven chromatin loop remodeling preferentially perturbs mature neuronal regulatory networks.

SV Del13 showed the strongest enrichment in MSN cells. Loop-associated DEGs were preferentially enriched for synaptic pathways, including SynGO synaptic processes, postsynaptic density, and presynaptic active zone gene sets (***Figure 4F***). Importantly, genes connected to altered loops were also enriched for schizophrenia associated loci identified in the most recent GWAS study (Trubetskoy et al. 2022) and brain-specific transcriptional programs, particularly cortical and excitatory neuron signatures, indicating that disruption of the TAD-b by SV Del13 preferentially affects neuronal regulatory networks involved in synaptic organization and disease-relevant transcriptional programs.

In contrast, SV Del10 exhibited its strongest enrichments in EGN cells, with comparatively weaker effects in MSN cells and minimal enrichment in NPCs. Loop-associated DEGs were preferentially enriched for chromatin regulatory programs involving CBX7, JARID2, MTF2, and RORC, together with cortical excitatory neuron signatures. Unlike SV Del13, SV Del10 also showed enrichment for genes implicated in neurodevelopmental disorders and genes under strong evolutionary constraint (LOEUF and pLI) but not specifically enriched for schizophrenia-associated signals ***(Figure 4G)***, suggesting that chromatin remodeling induced by this intradomain deletion preferentially affects developmental regulatory programs rather than synaptic networks.

## Discussion

We hypothesized that ultra-rare schizophrenia-associated SVs located at or near TAD-bs disrupt gene regulation through alterations in 3D genome organization and that these effects would be cell type-specific. To test our hypothesis, we performed an integrated analysis of Hi-C and transcriptomic data in NPCs, MSNs, and EGNs carrying chromosome 10 or chromosome 13 deletions identified in schizophrenia patients (Halvorsen et al. 2020). Our results demonstrate that the transcriptional consequences of these SVs are highly dependent on cellular context and are associated primarily with chromatin loop rewiring rather than widespread disruption of higher-order chromatin architecture.

Despite occurring within genomic regions predicted to contribute to domain organization, both deletions produced only modest alterations in higher-order chromatin architecture, including A/B compartments, TAD structure, and insulation profiles (***Figures 1 and 2***). Although local reductions in insulation were detected near the deleted regions, overall domain organization remained largely preserved across all three cell types (***Figure 2***). These observations are consistent with growing evidence that TAD-bs and large-scale chromosome architecture are relatively robust to structural perturbation, and that disruption of domain structure alone is often insufficient to induce widespread transcriptional dysregulation (Kabirova et al. 2024). In contrast, both deletions triggered extensive rewiring of chromatin loops, with hundreds of gained and lost interactions detected across cell types (***Figure 2C***). One possible interpretation is that structural variation exerts its effects primarily through remodeling of chromatin communication networks rather than through disruption of global chromosome organization, a model consistent with previous studies of disease-associated structural variation (Lupiáñez et al. 2015).

Remarkably, extensive loop remodeling translated into relatively limited direct transcriptional effects. Only a small fraction of differentially expressed genes overlapped altered chromatin interactions, and only a subset of rewired loops was associated with detectable changes in nearby gene expression (***Figure 4D*, *E***). These observations support emerging models in which chromatin architecture contributes to gene regulation in a probabilistic rather than deterministic manner, whereby multiple regulatory interactions can buffer the effects of individual chromatin contact changes (Sreenivasan et al. 2025). Thus, although chromatin looping provides an important framework for regulatory communication, alterations in individual contacts do not necessarily translate into immediate transcriptional consequences.

A notable finding was the predominance of enhancer-enhancer interactions among altered loops associated with differential gene expression (***Figure 4B*, *C***). While promoter-enhancer contacts have traditionally been viewed as the principal mechanism linking chromatin architecture to transcriptional regulation, increasing evidence suggests that enhancers often act collectively within higher-order chromatin hubs, where interactions among multiple regulatory elements contribute to gene regulation (Cornell et al. 2026; Oudelaar et al. 2018). Our findings therefore raise the possibility that these structural variants influence transcription not only through direct disruption of promoter regulation but also through perturbation of broader enhancer networks that coordinate neuronal gene expression programs.

The impact of chromatin loop rewiring differed substantially across developmental stages and neuronal lineages. NPCs exhibited relatively modest coupling between chromatin remodeling and differential gene expression, whereas both MSN and EGN populations displayed substantially greater loop rewiring and transcriptional dysregulation (***Figure 4D*, *E***). Developmental chromatin context may represent a critical determinant of SV penetrance, as the increasing complexity of enhancer landscapes and long-range regulatory interactions during neuronal differentiation could amplify the transcriptional consequences of structural variation (Kabirova et al. 2024; Rajarajan et al. 2018).

Importantly, the downstream consequences of chromatin rewiring differed between neuronal subtypes. MSNs showed particularly strong enrichment of CELF4-, FMRP-, and RBFOX-regulated gene networks (***Figure 4F*, G**), suggesting that structural variant-induced loop remodeling may preferentially perturb post-transcriptional regulatory programs in striatal neurons (Salamon et al. 2023; Wagnon et al. 2012; Gazzara et al. 2017; Weyn-Vanhentenryck et al. 2014). In contrast, EGNs exhibited stronger enrichment for chromatin remodeling pathways, including JARID2-, CHD7-, and CBX7-associated regulatory programs (Corley and Kroll 2015; Feng et al. 2017). These observations suggest that although the two deletions induce distinct patterns of local chromatin remodeling, their transcriptional consequences are strongly influenced by the regulatory architecture of each neuronal subtype, resulting in cell type-specific downstream molecular programs

Despite the lineage-specific differences in the magnitude of transcriptional responses and the specific genes affected across cell types, loop-associated DEGs were consistently enriched for chromatin regulatory processes. Rather than affecting isolated downstream genes, chromatin loop rewiring appears to preferentially perturb regulatory factors that occupy central positions within transcriptional and epigenetic networks. This convergence suggests that distinct structural variants may influence common neurodevelopmental regulatory programs despite acting through different local genomic mechanisms. Such convergence is consistent with genetic studies indicating that schizophrenia risk is highly heterogeneous at the variant level but tends to aggregate within a more restricted set of biological pathways governing neurodevelopment, synaptic function, RNA regulation, and gene expression control (Trubetskoy et al. 2022; Stauffer et al. 2023; Sullivan et al. 2024). Because many of the affected genes function as upstream regulatory factors, perturbation of a relatively small number of primary targets may propagate broadly through neuronal regulatory circuits, generating extensive secondary transcriptional responses. This framework provides a plausible explanation for why widespread differential gene expression was observed despite the relatively modest overlap between altered chromatin interactions and DEGs (Quinodoz et al. 2018; Di Giammartino et al. 2020; Oudelaar et al. 2018).

This study has several limitations. First, we modeled only two schizophrenia-associated TAD boundary-disrupting structural variants, and therefore our findings may not fully capture the diversity of mechanisms through which structural variants alter 3D genome organization. Second, although iPSC-derived neuronal models recapitulate key aspects of human neurodevelopment, they do not fully reflect the cellular diversity, circuit complexity, or maturation state of the human brain. Our analyses were also limited to three neuronal cell types and a single differentiation time point, precluding assessment of temporal changes in chromatin architecture during neuronal maturation. Integration of Hi-C and RNA-seq identifies candidate regulatory relationships but does not establish causal effects of chromatin looping on gene expression. Loop-associated genes were assigned based on the proximity of loop anchors (±5 kb) to gene coordinates, providing putative regulatory targets; however, the resolution of Hi-C precludes the precise assignment of individual regulatory elements, and long-range interactions may involve multiple target genes. Finally, because HICCUPSDiff is optimized to detect high-confidence focal loops, more subtle or diffuse changes in chromatin architecture may not have been captured, suggesting that our analyses likely provide a conservative estimate of the transcriptional consequences of chromatin remodeling induced by these structural variants.

Together, our findings support a model in which these two rare schizophrenia-associated structural variants can influence gene regulation through localized changes in chromatin connectivity, with effects shaped by cellular context. Despite their distinct genomic locations and mechanisms, these variants can converge on shared regulatory programs, highlighting three-dimensional genome organization as an important layer for understanding how structural variation contributes to schizophrenia risk.

## URLs

https://www.ncbi.nlm.nih.gov/geoprofiles

https://github.com/aidenlab/juicer

https://github.com/ArimaGenomics/mapping_pipeline

https://github.com/aidenlab/hic-straw

https://screen.encodeproject.org/

## Acknowledgements

This project was funded by NIMH R01 MH106611 (to J.P.S.) and NIMH MH108894 (to M.F.). We thank Dr. Adriana Beltrán and her team at the Human Pluripotent Cell Core Facility at the University of North Carolina at Chapel Hill for their assistance with the culturing and differentiation of iPSC for downstream genomics assays. High throughput sequencing was performed by the UNC Integrated Genomics Cores (RRID SCR_022620), which is supported in part by the School of Medicine Office of Research and funding from a Lineberger Cancer Center Support Grant (3P30CA016086-49S1) and the University Cancer Research Fund. We thank Dr Hemant Kelkar at the Bioinformatics and Analytics Research Collaborative at the University of North Carolina at Chapel Hill for the assistance with the bioinformatic analyses of the whole genome sequencing data.

## References

1. Amberger JS, Bocchini CA, Scott AF, Hamosh A. 2019. OMIM.org: leveraging knowledge across phenotype-gene relationships. Nucleic Acids Res 47: D1038–D1043.

2. Bryois J, Garrett ME, Song L, Safi A, Giusti-Rodriguez P, Johnson GD, Shieh AW, Buil A, Fullard JF, Roussos P, et al. 2018. Evaluation of chromatin accessibility in prefrontal cortex of individuals with schizophrenia. Nat Commun 9: 3121.

3. Bryois J, Skene NG, Hansen TF, Kogelman LJA, Watson HJ, Liu Z, Eating Disorders Working Group of the Psychiatric Genomics Consortium, International Headache Genetics Consortium, 23andMe Research Team, Brueggeman L, et al. 2020. Genetic identification of cell types underlying brain complex traits yields insights into the etiology of Parkinson’s disease. Nat Genet 52: 482–493.

4. Castro-Mondragon JA, Riudavets-Puig R, Rauluseviciute I, Lemma RB, Turchi L, Blanc-Mathieu R, Lucas J, Boddie P, Khan A, Manosalva Pérez N, et al. 2022. JASPAR 2022: the 9th release of the open-access database of transcription factor binding profiles. Nucleic Acids Res 50: D165–D173.

5. Chakraborty A, Wang JG, Ay F. 2022. dcHiC detects differential compartments across multiple Hi-C datasets. Nat Commun 13: 6827.

6. Chen M, Ju CJ-T, Zhou G, Chen X, Zhang T, Chang K-W, Zaniolo C, Wang W. 2019. Multifaceted protein-protein interaction prediction based on Siamese residual RCNN. Bioinformatics 35: i305–i314.

7. Chen R, Liu Y, Djekidel MN, Chen W, Bhattacherjee A, Chen Z, Scolnick E, Zhang Y. 2022. Cell type-specific mechanism of Setd1a heterozygosity in schizophrenia pathogenesis. Sci Adv 8: eabm1077.

8. Corley M, Kroll KL. 2015. The roles and regulation of Polycomb complexes in neural development. Cell Tissue Res 359: 65–85.

9. Cornell LJ, Harrold CL, Holliman S, Tsang F, Gosden ME, Hanssen LLP, Stolper R, Downes DJ, Biggs D, Preece C, et al. 2026. A functional overlap between actively transcribed genes and chromatin insulator elements. EMBO J 45: 2587–2613.

10. DeGroat W, Inoue F, Ashuach T, Yosef N, Ahituv N, Kreimer A. 2024. Comprehensive network modeling approaches unravel dynamic enhancer-promoter interactions across neural differentiation. Genome Biol 25: 221.

11. Di Giammartino DC, Polyzos A, Apostolou E. 2020. Transcription factors: building hubs in the 3D space. Cell Cycle 19: 2395–2410.

12. Dobin A, Davis CA, Schlesinger F, Drenkow J, Zaleski C, Jha S, Batut P, Chaisson M, Gingeras TR. 2013. STAR: ultrafast universal RNA-seq aligner. Bioinformatics 29: 15–21.

13. Durand NC, Robinson JT, Shamim MS, Machol I, Mesirov JP, Lander ES, Aiden EL. 2016a. Juicebox Provides a Visualization System for Hi-C Contact Maps with Unlimited Zoom. Cell Syst 3: 99–101.

14. Durand NC, Shamim MS, Machol I, Rao SSP, Huntley MH, Lander ES, Aiden EL. 2016b. Juicer Provides a One-Click System for Analyzing Loop-Resolution Hi-C Experiments. Cell Syst 3: 95–98.

15. ENCODE Project Consortium, Moore JE, Purcaro MJ, Pratt HE, Epstein CB, Shoresh N, Adrian J, Kawli T, Davis CA, Dobin A, et al. 2020. Expanded encyclopaedias of DNA elements in the human and mouse genomes. Nature 583: 699–710.

16. Espuny-Camacho I, Michelsen KA, Gall D, Linaro D, Hasche A, Bonnefont J, Bali C, Orduz D, Bilheu A, Herpoel A, et al. 2013. Pyramidal neurons derived from human pluripotent stem cells integrate efficiently into mouse brain circuits in vivo. Neuron 77: 440–456.

17. Ewels P, Magnusson M, Lundin S, Käller M. 2016. MultiQC: summarize analysis results for multiple tools and samples in a single report. Bioinformatics 32: 3047–3048.

18. Feng W, Kawauchi D, Körkel-Qu H, Deng H, Serger E, Sieber L, Lieberman JA, Jimeno-González S, Lambo S, Hanna BS, et al. 2017. Chd7 is indispensable for mammalian brain development through activation of a neuronal differentiation programme. Nat Commun 8: 14758.

19. Gazzara MR, Mallory MJ, Roytenberg R, Lindberg JP, Jha A, Lynch KW, Barash Y. 2017. Ancient antagonism between CELF and RBFOX families tunes mRNA splicing outcomes. Genome Res 27: 1360–1370.

20. The Gene Ontology Consortium. 2019. The Gene Ontology Resource: 20 years and still GOing strong. Nucleic Acids Res 47: D330–D338.

21. Girdhar K, Hoffman GE, Bendl J, Rahman S, Dong P, Liao W, Hauberg ME, Sloofman L, Brown L, Devillers O, et al. 2022. Chromatin domain alterations linked to 3D genome organization in a large cohort of schizophrenia and bipolar disorder brains. Nat Neurosci 25: 474–483.

22. Giusti-Rodríguez P, Lu L, Yang Y, Crowley CA, Liu X, Juric I, Martin JS, Abnousi A, Allred SC, Ancalade N, et al. 2019. Using three-dimensional regulatory chromatin interactions from adult and fetal cortex to interpret genetic results for psychiatric disorders and cognitive traits. 406330. https://www.biorxiv.org/content/10.1101/406330v2 (Accessed September 8, 2026).

23. GTEx Consortium. 2020. The GTEx Consortium atlas of genetic regulatory effects across human tissues. Science 369: 1318–1330.

24. Halvorsen M, Huh R, Oskolkov N, Wen J, Netotea S, Giusti-Rodriguez P, Karlsson R, Bryois J, Nystedt B, Ameur A, et al. 2020. Increased burden of ultra-rare structural variants localizing to boundaries of topologically associated domains in schizophrenia. Nat Commun 11: 1842.

25. Hu B, Won H, Mah W, Park RB, Kassim B, Spiess K, Kozlenkov A, Crowley CA, Pochareddy S, PsychENCODE Consortium, et al. 2021. Neuronal and glial 3D chromatin architecture informs the cellular etiology of brain disorders. Nat Commun 12: 3968.

26. Hua D, Gu M, Zhang X, Du Y, Xie H, Qi L, Du X, Bai Z, Zhu X, Tian D. 2024. DiffDomain enables identification of structurally reorganized topologically associating domains. Nat Commun 15: 502.

27. Jacobson M. 1985. Clonal analysis and cell lineages of the vertebrate central nervous system. Annu Rev Neurosci 8: 71–102.

28. Kabirova E, Ryzhkova A, Lukyanchikova V, Khabarova A, Korablev A, Shnaider T, Nuriddinov M, Belokopytova P, Smirnov A, Khotskin NV, et al. 2024. TAD border deletion at the Kit locus causes tissue-specific ectopic activation of a neighboring gene. Nat Commun 15: 4521.

29. Karczewski KJ, Francioli LC, Tiao G, Cummings BB, Alföldi J, Wang Q, Collins RL, Laricchia KM, Ganna A, Birnbaum DP, et al. 2020. The mutational constraint spectrum quantified from variation in 141,456 humans. Nature 581: 434–443.

30. Koesterich J, Liu J, Williams SE, Yang N, Kreimer A. 2024. Network Analysis of Enhancer-Promoter Interactions Highlights Cell-Type-Specific Mechanisms of Transcriptional Regulation Variation. Int J Mol Sci 25: 9840.

31. Koopmans F, van Nierop P, Andres-Alonso M, Byrnes A, Cijsouw T, Coba MP, Cornelisse LN, Farrell RJ, Goldschmidt HL, Howrigan DP, et al. 2019. SynGO: An Evidence-Based, Expert-Curated Knowledge Base for the Synapse. Neuron 103: 217–234.e4.

32. Kruse K, Hug CB, Vaquerizas JM. 2020. FAN-C: a feature-rich framework for the analysis and visualisation of chromosome conformation capture data. Genome Biol 21: 303.

33. Lek M, Karczewski KJ, Minikel EV, Samocha KE, Banks E, Fennell T, O’Donnell-Luria AH, Ware JS, Hill AJ, Cummings BB, et al. 2016. Analysis of protein-coding genetic variation in 60,706 humans. Nature 536: 285–291.

34. Li C, Bonder MJ, Syed S, Jensen M, Human Genome Structural Variation Consortium (HGSVC), HGSVC Functional Analysis Working Group, Gerstein MB, Zody MC, Chaisson MJP, Talkowski ME, et al. 2024. An integrative TAD catalog in lymphoblastoid cell lines discloses the functional impact of deletions and insertions in human genomes. Genome Res 34: 2304–2318.

35. Li H. 2013. Aligning sequence reads, clone sequences and assembly contigs with BWA-MEM. http://arxiv.org/abs/1303.3997 (Accessed September 8, 2026).

36. Li H, Durbin R. 2010. Fast and accurate long-read alignment with Burrows-Wheeler transform. Bioinformatics 26: 589–595.

37. Li H, Durbin R. 2009. Fast and accurate short read alignment with Burrows-Wheeler transform. Bioinformatics 25: 1754–1760.

38. Lieberman-Aiden E, van Berkum NL, Williams L, Imakaev M, Ragoczy T, Telling A, Amit I, Lajoie BR, Sabo PJ, Dorschner MO, et al. 2009. Comprehensive mapping of long-range interactions reveals folding principles of the human genome. Science 326: 289–293.

39. Lin C, Jain S, Kim H, Bar-Joseph Z. 2017. Using neural networks for reducing the dimensions of single-cell RNA-Seq data. Nucleic Acids Res 45: e156.

40. Love MI, Huber W, Anders S. 2014. Moderated estimation of fold change and dispersion for RNA-seq data with DESeq2. Genome Biol 15: 550.

41. Lu L, Liu X, Huang W-K, Giusti-Rodríguez P, Cui J, Zhang S, Xu W, Wen Z, Ma S, Rosen JD, et al. 2020. Robust Hi-C Maps of Enhancer-Promoter Interactions Reveal the Function of Non-coding Genome in Neural Development and Diseases. Mol Cell 79: 521–534.e15.

42. Lupiáñez DG, Kraft K, Heinrich V, Krawitz P, Brancati F, Klopocki E, Horn D, Kayserili H, Opitz JM, Laxova R, et al. 2015. Disruptions of topological chromatin domains cause pathogenic rewiring of gene-enhancer interactions. Cell 161: 1012–1025.

43. Marshall CR, Howrigan DP, Merico D, Thiruvahindrapuram B, Wu W, Greer DS, Antaki D, Shetty A, Holmans PA, Pinto D, et al. 2017. Contribution of copy number variants to schizophrenia from a genome-wide study of 41,321 subjects. Nat Genet 49: 27–35.

44. Maury EA, Sherman MA, Genovese G, Gilgenast TG, Kamath T, Burris SJ, Rajarajan P, Flaherty E, Akbarian S, Chess A, et al. 2023. Schizophrenia-associated somatic copy-number variants from 12,834 cases reveal recurrent NRXN1 and ABCB11 disruptions. Cell Genom 3: 100356.

45. McKenna A, Hanna M, Banks E, Sivachenko A, Cibulskis K, Kernytsky A, Garimella K, Altshuler D, Gabriel S, Daly M, et al. 2010. The Genome Analysis Toolkit: a MapReduce framework for analyzing next-generation DNA sequencing data. Genome Res 20: 1297–1303.

46. Meeker ND, Hutchinson SA, Ho L, Trede NS. 2007. Method for isolation of PCR-ready genomic DNA from zebrafish tissues. Biotechniques 43: 610, 612, 614.

47. Open2C, Abdennur N, Abraham S, Fudenberg G, Flyamer IM, Galitsyna AA, Goloborodko A, Imakaev M, Oksuz BA, Venev SV, et al. 2024. Cooltools: Enabling high-resolution Hi-C analysis in Python. PLoS Comput Biol 20: e1012067.

48. Oudelaar AM, Davies JOJ, Hanssen LLP, Telenius JM, Schwessinger R, Liu Y, Brown JM, Downes DJ, Chiariello AM, Bianco S, et al. 2018. Single-allele chromatin interactions identify regulatory hubs in dynamic compartmentalized domains. Nat Genet 50: 1744–1751.

49. Owen MJ, Legge SE, Rees E, Walters JTR, O’Donovan MC. 2023. Genomic findings in schizophrenia and their implications. Mol Psychiatry 28: 3638–3647.

50. Patro R, Duggal G, Love MI, Irizarry RA, Kingsford C. 2017. Salmon provides fast and bias-aware quantification of transcript expression. Nat Methods 14: 417–419.

51. Quinlan AR, Hall IM. 2010. BEDTools: a flexible suite of utilities for comparing genomic features. Bioinformatics 26: 841–842.

52. Quinodoz SA, Ollikainen N, Tabak B, Palla A, Schmidt JM, Detmar E, Lai MM, Shishkin AA, Bhat P, Takei Y, et al. 2018. Higher-Order Inter-chromosomal Hubs Shape 3D Genome Organization in the Nucleus. Cell 174: 744–757.e24.

53. Rajarajan P, Borrman T, Liao W, Schrode N, Flaherty E, Casiño C, Powell S, Yashaswini C, LaMarca EA, Kassim B, et al. 2018. Neuron-specific signatures in the chromosomal connectome associated with schizophrenia risk. Science 362: eaat4311.

54. Rao SSP, Huntley MH, Durand NC, Stamenova EK, Bochkov ID, Robinson JT, Sanborn AL, Machol I, Omer AD, Lander ES, et al. 2014. A 3D map of the human genome at kilobase resolution reveals principles of chromatin looping. Cell 159: 1665–1680.

55. Robinson JT, Thorvaldsdottir H, Turner D, Mesirov JP. 2023. igv.js: an embeddable JavaScript implementation of the Integrative Genomics Viewer (IGV). Bioinformatics 39: btac830.

56. Robinson JT, Thorvaldsdóttir H, Wenger AM, Zehir A, Mesirov JP. 2017. Variant Review with the Integrative Genomics Viewer. Cancer Res 77: e31–e34.

57. Robinson JT, Thorvaldsdóttir H, Winckler W, Guttman M, Lander ES, Getz G, Mesirov JP. 2011a. Integrative genomics viewer. Nat Biotechnol 29: 24–26.

58. Robinson JT, Thorvaldsdóttir H, Winckler W, Guttman M, Lander ES, Getz G, Mesirov JP. 2011b. Integrative genomics viewer. Nat Biotechnol 29: 24–26.

59. Salamon I, Park Y, Miškić T, Kopić J, Matteson P, Page NF, Roque A, McAuliffe GW, Favate J, Garcia-Forn M, et al. 2023. Celf4 controls mRNA translation underlying synaptic development in the prenatal mammalian neocortex. Nat Commun 14: 6025.

60. Sreenivasan VKA, Yumiceba V, Spielmann M. 2025. Structural variants in the 3D genome as drivers of disease. Nat Rev Genet 26: 742–760.

61. Stauffer E-M, Bethlehem RAI, Dorfschmidt L, Won H, Warrier V, Bullmore ET. 2023. The genetic relationships between brain structure and schizophrenia. Nat Commun 14: 7820.

62. Sullivan PF, Meadows JRS, Gazal S, Phan BN, Li X, Genereux DP, Dong MX, Bianchi M, Andrews G, Sakthikumar S, et al. 2023. Leveraging Base Pair Mammalian Constraint to Understand Genetic Variation and Human Disease.

63. Sullivan PF, Yao S, Hjerling-Leffler J. 2024. Schizophrenia genomics: genetic complexity and functional insights. Nat Rev Neurosci 25: 611–624.

64. Tam RW, Keung AJ. 2022. Human Pluripotent Stem Cell-Derived Medium Spiny Neuron-like Cells Exhibit Gene Desensitization. Cells 11: 1411.

65. Thorvaldsdóttir H, Robinson JT, Mesirov JP. 2013. Integrative Genomics Viewer (IGV): high-performance genomics data visualization and exploration. Brief Bioinform 14: 178–192.

66. Trubetskoy V, Pardiñas AF, Qi T, Panagiotaropoulou G, Awasthi S, Bigdeli TB, Bryois J, Chen C-Y, Dennison CA, Hall LS, et al. 2022. Mapping genomic loci implicates genes and synaptic biology in schizophrenia. Nature 604: 502–508.

67. Visel A, Minovitsky S, Dubchak I, Pennacchio LA. 2007. VISTA Enhancer Browser--a database of tissue-specific human enhancers. Nucleic Acids Res 35: D88–92.

68. Wagnon JL, Briese M, Sun W, Mahaffey CL, Curk T, Rot G, Ule J, Frankel WN. 2012. CELF4 regulates translation and local abundance of a vast set of mRNAs, including genes associated with regulation of synaptic function. PLoS Genet 8: e1003067.

69. Wang D, Liu S, Warrell J, Won H, Shi X, Navarro FCP, Clarke D, Gu M, Emani P, Yang YT, et al. 2018. Comprehensive functional genomic resource and integrative model for the human brain. Science 362: eaat8464.

70. Weyn-Vanhentenryck SM, Mele A, Yan Q, Sun S, Farny N, Zhang Z, Xue C, Herre M, Silver PA, Zhang MQ, et al. 2014. HITS-CLIP and integrative modeling define the Rbfox splicing-regulatory network linked to brain development and autism. Cell Rep 6: 1139–1152.

71. Yang A, Chen J, Zhao X-M. 2021. nMAGMA: a network-enhanced method for inferring risk genes from GWAS summary statistics and its application to schizophrenia. Brief Bioinform 22: bbaa298.

72. Babraham Bioinformatics - FastQC A Quality Control tool for High Throughput Sequence Data. https://www.bioinformatics.babraham.ac.uk/projects/fastqc/.

